# Somatic HCN channels augment and speed up GABAergic basket cell input-output function in human neocortex

**DOI:** 10.1101/2021.06.09.447671

**Authors:** Viktor Szegedi, Emőke Bakos, Szabina Furdan, Pal Barzo, Gabor Tamas, Karri Lamsa

## Abstract

Neurons in the mammalian brain exhibit evolution-driven species-specific differences in their functional properties. Therefore, understanding the human brain requires unraveling the human neuron “uniqueness” and how it contributes to the operation of specific neuronal circuits. We show here that a highly abundant type of inhibitory neurons in the neocortex, GABAergic parvalbumin-expressing basket cell (pv+BC), exhibits in the human brain a specific somatic leak current mechanism, which is absent in their rodent neuronal counterparts. Human pv+BC soma shows electric leak conductance mediated by hyperpolarization-activated cyclic nucleotide-gated channels. This leak conductance has depolarizing effects on the resting membrane potential and it accelerates the rise of synaptic potentials in the cell soma. The leak facilitates the human pv+BC input-to-output fidelity and shortens the action potential generation to excitatory inputs. This mechanism constitutes an adaptation that enhances signal transmission fidelity and speed in the common inhibitory circuit in the human but not in the rodent neocortex.

## INTRODUCTION

Fast-spiking GABAergic interneurons of the mammalian cerebral cortex play central role in cognitive and sensory processes by supplying accurately timed inhibition of neuronal networks ^1, 2^. Precise and faithful operation of these parvalbumin-expressing (pv+) neurons relies on “fast in-fast out” function, by which they transform excitatory synaptic inputs to inhibitory outputs with a high fidelity, short latency and sub-millisecond temporal precision (see for instance ^3^). Time-lag and jitter in their operation mostly arise from the transformation of excitatory postsynaptic potentials (EPSPs) into a spiking output ^4, 5^. This EPSP-spike transformation takes place in (or very close to) cell soma and the process is well-documented in rodent cortical neurons including the pv+ GABAergic cell population ^1, 6, 7^. Yet very few studies have investigated the EPSP-spike transformation or the input-to-output function in human neurons^8, 9, 10, 11, 12, 13^ and particularly the GABAergic human neuronal types have received little attention. Understanding the functioning of human neurons is highly relevant, because findings in experimental animals do not always translate to human. In fact, analogous neuronal types between the human and rodent cortex exhibit various between-species differences in their physiological functional parameters ^9, 13, 14, 15, 16, 17, 18, 19, 20, 21^. Species-specific behaviors may hence arise from even small differences in neuronal types and neuronal circuits ^22, 23^, making it critical to uncover the “uniqueness” of the human neurons and to investigate its outcome on the operation of anatomically identified neuronal circuits.

Although single-cell RNA sequencing studies denote relatively conserved gene expression patterns among GABAergic inhibitory neurons of the mammalian neocortex ^24, 25^, analogous neuronal types such as pv+ GABAergic interneurons show inter-species differences in terms of neurite arborization ^26^, ion channel expression ^27^, as well as temporal dimensions of basal synaptic transmission ^14, 28^. In general, pv+ cortical interneurons are characterized by rapid action potentials and a high-frequency firing capacity across various experimental species ^10, 29, 30, 31, 32^, but action potential firing and intrinsic electrical properties of the cells differ quantitatively between species ^28, 31, 33^. However, it remains unknown how these affect EPSP-spike transformation and input-to-output function of the most numerous neocortical inhibitory interneuronal type, the pv+ basket cell (BC) ^34, 35^ in human.

Excitability in cell soma in many neurons is enhanced by hyperpolarization-activated cyclic nucleotide-gated (HCN) channels and these provide a well-documented mechanism allowing to facilitate EPSP kinetics and the EPSP-spike coupling ^33, 36, 37, 38, 39^. HCN channels are voltage-gated cation channels opening at potentials negative to −50 mV that regulate resting membrane potential and intrinsic excitability by their permeability to K^+^ and Na^+^ ions, with a reversal potential of approximately −30 mV ^40, 41^. RNA sequencing studies and protein analyses indicate high levels of HCN-channel expression in the human and rodent neocortex ^18, 42^. In neocortical pv+ GABAergic neurons, which contain primarily the HCN-channel type 1 (HCN1) subunit ^40, 43, 44, 45^, rodent experiments showed their absence in the soma. Although pv+ neurons in some other areas of brain exhibit somatic HCN-channel activity ^40, 46, 47^, in the neocortex the HCN channels are localized in their axon ^42, 48, 49^. This is obvious in many studies demonstrating minuscule hyperpolarization-evoked voltage sag potential, which is a hallmark of somatic HCN activity, in neocortical pv+ cells. Furthermore, HCN-channel blockers barely have any effect on the rodent neocortical pv+ cell resting membrane potential or somatic excitability ^33, 36, 37, 38, 39, 43, 50^. Nevertheless, HCN-channel activity and its localization in human pv+ neurons have remained elusive.

We investigated somatic HCN-channel activity and its role in the transformation of excitatory inputs into a spike in human pv+BCs among neocortical layers 2/3 (L2/3) where neurons make major contributions to corticocortical connections and function to integrate information among cortical areas (for instance see ^51^). The study was performed using acute human brain slices from non-pathological neocortical tissue resected in deep area brain surgery. We report robust HCN-channel activity in the somatic compartment of human, but not mouse, anatomically-identified parvalbumin-immunopositive BCs. Somatic HCN conductance depolarizes the resting potential, reduces excitatory synaptic potential (EPSP) onset-to-peak time, and shortens the delay from excitatory somatic currents to spike in human interneurons. These findings indicate that somatic HCN-channel activity promotes signal transmission through this anatomically-identified inhibitory neuronal circuit in the human neocortex. We suggest that this mechanism represents an evolutionary adaptation to increase fidelity, speed, and precision of signal transmission through this common inhibitory circuit in the human neocortex.

## RESULTS

We studied 45 pv+BCs in the human neocortex L2/3 to assess HCN-channel mediated activity in their soma. Cells were recorded in whole-cell clamp within acute neocortical slices prepared from tissue material that had to be removed in surgery to gain access to deeper-brain targets. Cells were visualized and identified by their axon forming boutons around unlabeled L2/3 neurons, and by positive immunoreaction for pv (pv+; n = 45). For comparison, we studied 38 visualized fast-spiking BCs in the mouse somatosensory cortex (Supplementary Table 1).

### pv+BCs show large HCN-channel-mediated somatic sag potential in human but not in mouse

We found that pv+BCs in the human neocortex show robust hyperpolarization-activated sag potential. First, we recorded cells in current clamp by applying square-pulse current steps (250 ms) to hyperpolarize or depolarize the membrane potential (range from −95 mV to +40 mV) at a holding potential of −70 mV (Fig. 1A1–2). The hyperpolarizing square-pulse steps evoked voltage sag during the step, and a rebound voltage sag of similar amplitude but opposite polarity was measured following the membrane potential step ^18^ (P = 0.516, Mann–Whitney U-test) (Supplementary Table 2). This property is characteristic of HCN-channel activation during hyperpolarization and of HCN-channel inactivation by repolarization, respectively ^18, 40^ (see Fig. 1A1–2). In addition, the HCN-channel antagonist ZD7288 (10 μM) similarly blocked the voltage sag and the rebound sag (Fig. 1A3).

**Figure 1.**
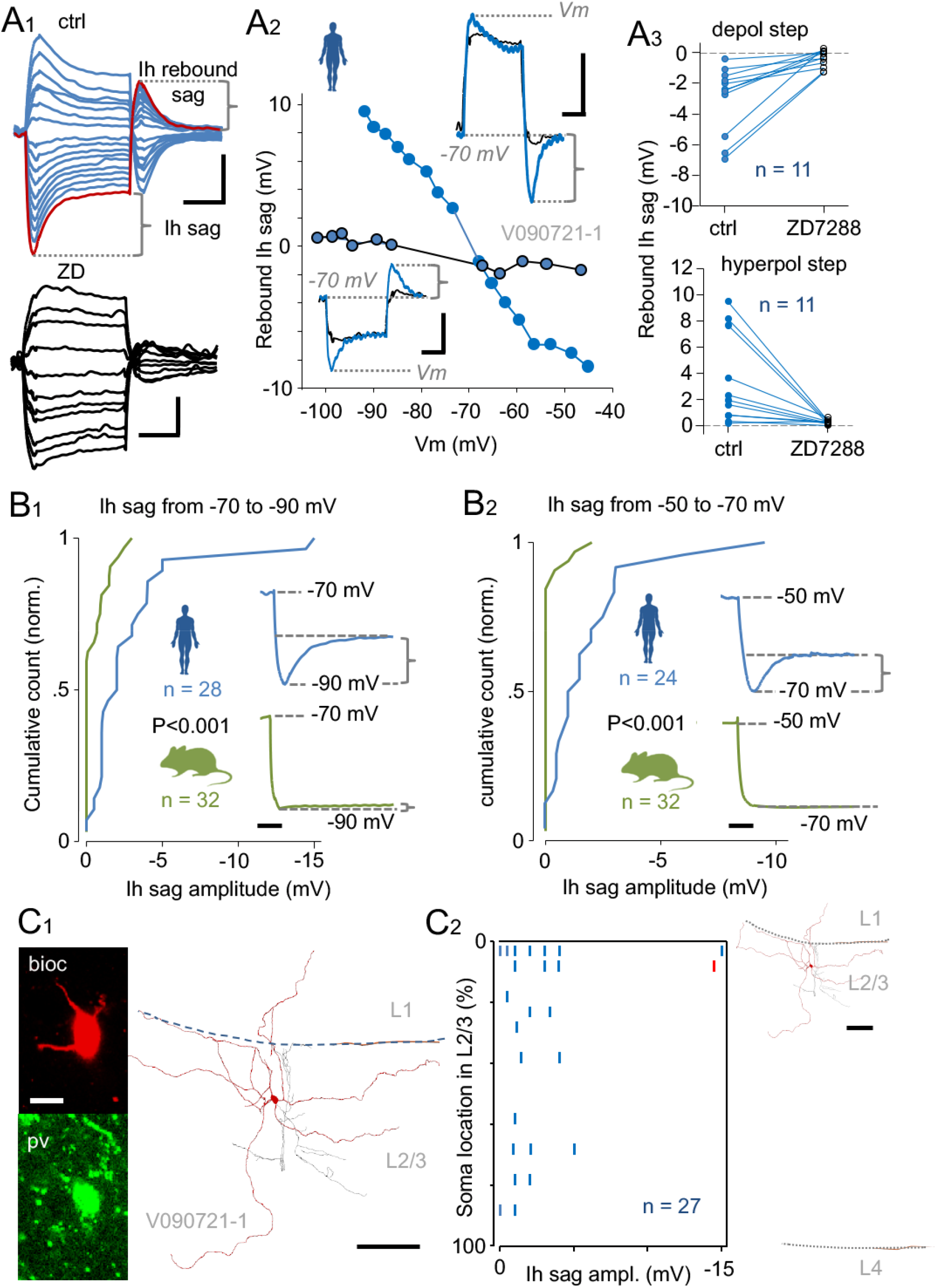
Human compared to mouse pv+BCs show a robust somatic HCN-channel-mediated sag potential. *A1*. Human pv+BC somatic membrane potential responses to depolarizing or hyperpolarizing square-pulse current steps delivered at −70 mV. Traces in control (blue) show characteristic hyperpolarization-activated sag potential and its blockade by ZD7288 (ZD; 10 μM) (black traces). One trace in control conditions is shown in red to illustrate the ZD-sensitive sag potential amplitude during a hyperpolarizing step, and rebound sag at resting membrane potential (Em) following the step (scale 10 mV, 100 ms). *A2*. Plot shows the rebound sag amplitude (ordinate) measured in a pv+BC at −70 mV following membrane potential steps (Vm, see insets) in the control conditions (blue symbols) and in the presence of ZD (black). The rebound sag amplitude gives a reliable measure that allows to compare the strength of HCN-channel mediated responses at different somatic membrane potentials, with the sag amplitude systematically measured at −70 mV to prevent any variable electrochemical driving force. Insets: Specimen membrane potential traces of a depolarizing or hyperpolarizing step in control (blue) and in ZD (black) conditions. HCN rebound sag amplitude is defined in traces with dotted lines and a bracket (scale 10 mV, 100 ms). *A3*. The rebound HCN sag amplitude measured in similar way in 11 human pv+BCs following depolarizing voltage steps (from −70 mV to −50 mV) and hyperpolarizing steps (to −90 mV from −70 mV). Blue shows control and black the ZD (10 μM) conditions (P < 0.001 and P < 0.001, Wilcoxon Signed-Rank Test). *B*. Larger somatic HCN sag potential observed in human compared to mouse pv+BCs. *B1*. The cumulative histogram shows HCN sag potential amplitude (as illustrated in one cell in inset) evoked by hyperpolarizing somatic step from −70 mV to −90 mV in identified pv+BCs. The HCN sag amplitude is larger in human cells (blue, n = 28) compared to mouse cells (green, n = 32) (P < 0.001, Mann–Whitney U-test). Insets: Traces illustrate robust HCN voltage sag in a human but not in a mouse pv+BC (scale 50 ms). *B2*. Cumulative histogram of HCN sag amplitude in human and mouse pv+BCs by membrane potential steps from −50 mV to −70 mV. The HCN sag amplitude in human is larger than in mouse (P < 0.001, Mann– Whitney U-test). Insets: Sample traces in a human and in a mouse pv+BCs (scale 10 mV, 50 ms). *C*. A robust HCN sag occurs in the human pv+BCs among the superficial or supragranular L2/3. *C1*. Illustration of one human pv+BC in L2/3. Confocal imaging micrographs show positive immunofluorescence reaction of the biocytin-filled (bioc) neuron to parvalbumin (pv) with a soma location at superficial L2. Left: Fluorescence micrographs illustrate biocytin (bioc)-filled soma immunopositive for parvalbumin (pv). Scale: 10 μm. Partial anatomical reconstruction shows cell soma and dendrites (red), as well as axon (black) in two 60 μm-thick merged sections. Scale: 100 μm. Horizontal lines shows L1 and L4 borders. *C2*. Abscissa summarizes HCN sag amplitude in human pv+BC and ordinate shows the soma location in L2/3 relative to L1 border (indicated as 0%) and to L4 border (indicated as 100% in the plot). Percentage bin is 5%. Red dot in the plot belongs to the sample cell shown on the right in full scale (scale 100 μm). Sag values are from 27 cells in *B1* data with a successful anatomical localization of the soma.

We next compared somatic sag robustness between human and mouse pv+BCs by using hyperpolarizing voltage step at −70 mV. In human (n = 28 cells), hyperpolarizing step to −90.6 mV (average) showed a 2.91 mV average sag potential amplitude, with a large range observed between cells from 0 to 15 mV. In mouse (n = 32 cells), a similar hyperpolarizing step (to −86.4 mV average) showed 0.53 mV sag, with values ranging from 0 to 3.0 mV (Fig. 1B1). A difference in sag amplitude between human and mouse cells was also seen when a hyperpolarizing membrane potential step was applied to −70 mV at −50 mV (Fig. 1B2).

Hyperpolarization of human pv+BCs from −49.3 mV (average) to −70.5 mV revealed a 1.9 mV average sag with a range from 0 to 9.5 mV. Correspondingly, in mouse pv+BCs, a step from −46.1 mV to −71.0 mV on average showed a barely detectable sag of just 0.2 mV on average with a range from 0 to 2 mV. Comparing between human and mouse pv+BCs further showed a statistically significant difference in somatic voltage sag amplitude at both step protocols (P < 0.001, Mann–Whitney U-test) (Fig. 1B1-2).

The pv+BCs were systematically filled with biocytin during recording and their anatomical location was determined *post hoc* (Fig. 1C1). We therefore examined the vertical position of pv+BCs soma in L2/3 to assess whether the voltage sag strength differed between cells localized in the superficial or deep L2/3 (Fig. 1C2). Yet we failed to find correlation between sag amplitude and the depth of cell soma in L2/3 (P = 0.298, correlation coefficient = 0.207, Spearman correlation). The soma location for each cell was measured as their relative positioning from the L1 border to the L4 border (see Fig. 1C2) (see Methods).

### HCN channels depolarize somatic resting membrane potential in human pv+BCs

We next studied somatic HCN-channel activity in 18 randomly selected human and in 18 randomly selected mouse pv+BCs that were kept at resting membrane potential (Em). By applying a robust depolarizing subthreshold step to −50 mV (250 ms, average in human = −48.1 mV, mouse = −46.0 mV, n = 18 and 18), we aimed to inactivate HCN channels, which are known to reactivate by repolarization to Em and to generate a voltage sag ^40, 42^. Figure 2A summarizes the results showing the Em of human and mouse pv+BCs (Fig. 2A1) and the voltage sag amplitude that was measured at Em in these cells following the depolarizing step (Fig. 2A2).

**Figure 2.**
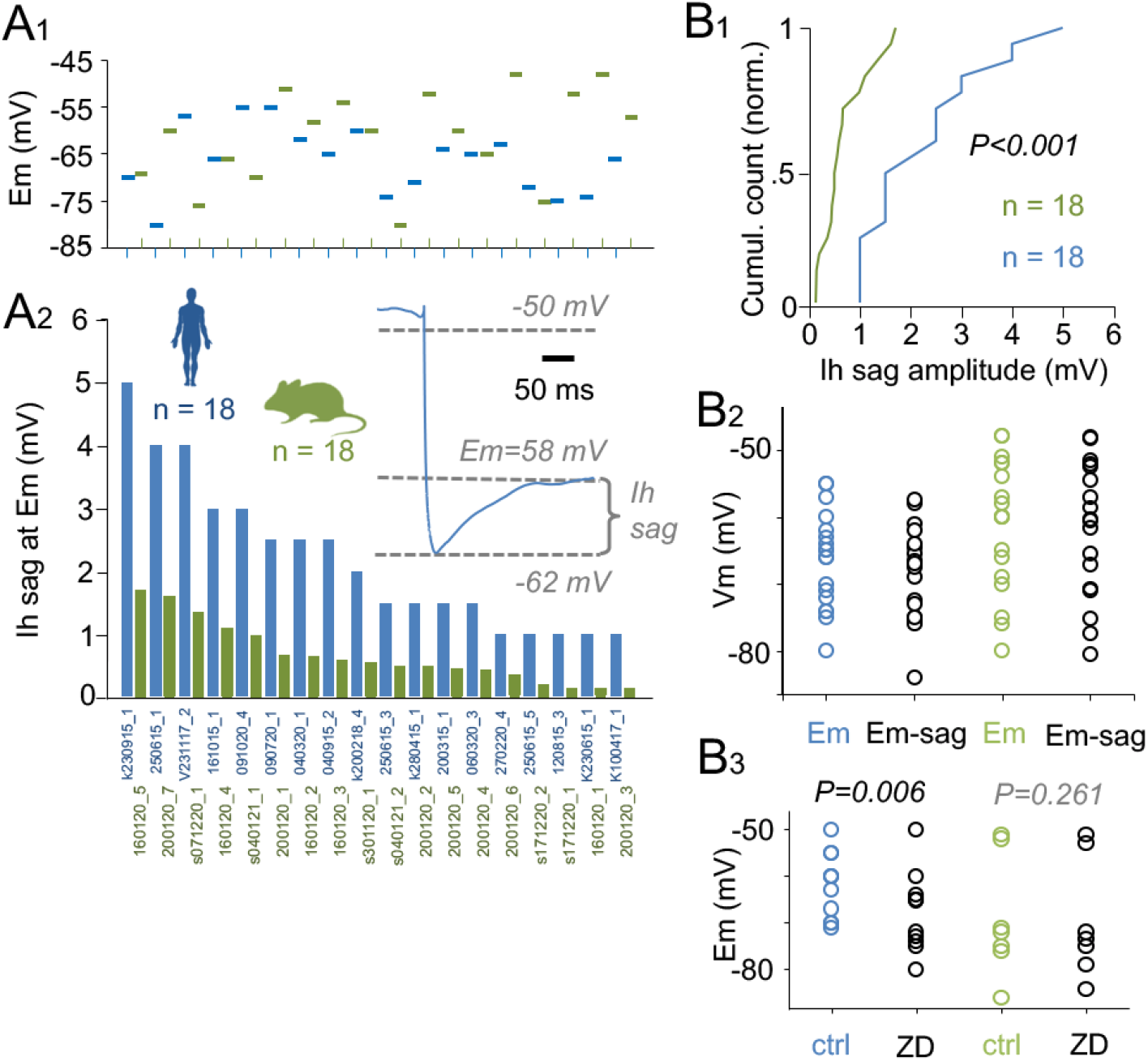
HCN-channel activity contributes to determining human pv+BC resting membrane potential. *A1*. Symbols show the resting membrane potential (Em) in 18 human and 18 mouse pv+BCs (blue and green, respectively) (Em between the species P = 0.080, Student’s t-test) (human −66.33 ± 7.15 mV, mean ± standard deviation (SD), n = 18; and mouse −61.17 ± 9.84 mV, n = 18). *A2*. HCN sag amplitude in the pv+BCs measured at Em following a depolarizing step to −50 mV. The robust depolarizing pulse aims to inactivate the HCN channels, which reactivate at Em by the repolarization as illustrated in the inset. Bar histogram shows HCN sag amplitude from the largest to the smallest in human and mouse pv+BCs. Individual cells are identified by codes in abscissa. *B1*. Cumulative histogram collates HCN sag amplitudes at Em in the human and mouse pv+BCs. The sag amplitude is larger in human than in mouse pv+BCs (P < 0.001, Mann– Whitney U-test). *B2*. Plot summarizes the Em values (Em) and Vm values in the cells when a depolarizing effect of the sag amplitude (when applicable) is subtracted (E-sag). *B3*. Dataset from small number of separate experiments show that the ZD wash-in generates similar negative shift of Em in human (n = 11) but not in mouse (n = 7) pv+BCs (paired Student’s t-test).

Altogether, human cells showed a 2.20 mV average sag potential amplitude with a range from 1.0 to 5.1 mV at their Em (average −66.3 mV, n = 18). For comparison, the identified pv+BCs in mouse showed a 0.66 mV average sag amplitude with a range from 0 to 1.7 mV at Em (average −61.2 mV, n = 18) (Fig. 1B1). Subtracting the voltage sag amplitude from the Em value measured at steady-state (see Fig. 2A2 inset) revealed the depolarizing effect of HCN-channel activity on their somatic membrane potential under resting conditions. Figure 2B2 sums up this depolarizing effect on Em in both human and mouse pv+BCs. However, somatic HCN activity fails to explain the diversity of Em values observed between individual pv+BCs (Supplementary Fig. S1).

In addition, the depolarizing effect of HCN-channel activity on Em was measured in other set of experiments, in which ZD7288 (ZD; 10 μM) was washed-in while monitoring Em in pv+BCs. In 11 human and 7 mouse pv+BCs, we measured Em just before ZD application and after 3–5 min in presence of the drug. We found a −5.65 mV average shift of Em (in control −61.9 ± 6.9 mV) in human pv+BCs (P = 0.006) compared to a −0.69 mV average change in mouse pv+BCs (n = 7, in control −69.0 ± 12.9 mV) (P = 0.261) (paired Student’s t-test). The Em data in control conditions and in the presence ZD are summarized in Figure 2B3. The relatively small sample size in these experiments is explained by fact that in most recordings with ZD, Em was not monitored during ZD application.

### HCN channels maintain somatic leak conductance in human pv+BCs at resting potential

In human pv+BCs, we obtained robust immunohistochemical staining for HCN1, the common channel isoform and the major HCN-channel protein in cortical neurons ^40, 43, 45^.

We performed double immunohistochemical labeling for pv and HCN1 in three human tissue samples (Fig. 3A). The immunohistochemical results show HCN1 in human pv+BCs. The results support our electrophysiological findings described above in current clamp showing HCN-channel activity in the somatic compartment in human pv+BCs. Indeed, we verified HCN channel-mediated electric conductance in human pv+BC soma. We performed voltage-clamp recording in five human and five mouse pv+BCs that were selected randomly for the experiments. We voltage-clamped the cells at their Em (average in human of −59.0 mV, mouse of −70.6 mV), and measured somatic leaking conductance (Gleak) by using brief voltage-clamp steps (−10 or 10 mV, 10 ms applied at 0.1 Hz) while washing-in the HCN-blocker ZD (10 μM). In human pv+BCs, Gleak was 3.78 nS, 2.97-4.46 nS (median, quartiles) in baseline control conditions and 3.32 nS, 2.31-3.88 nS following application of ZD (3–5 min). Thus, ZD reduced Gleak to 0.84 (average, range 0.57 to 1.04) from baseline control. In mouse pv+BCs, Gleak was 4.87 nS, 4.75-6.17 nS in baseline control conditions. In the presence of ZD, baseline-normalized Gleak was 0.96 (average, range 0.94 to 0.98). The ZD-sensitive conductance generated through HCN channels (G_HCN_) was of 0.58 nS (average range of 0–1.42 nS) in the human pv+BCs, and of 0.21 nS (average range of 0.12–0.28 nS) in the mouse pv+BCs. Figure 3B and C illustrate the conductance measurement made in these experiments. Figure 3D summarizes the measured leaking conductance in all pv+BCs in control conditions and in the presence of ZD. Leaking conductance was measured from the voltage step current at the end of step when it approached a steady-state level, while the amplitude is in a linear relation to the soma leaking conductance.

**Figure 3.**
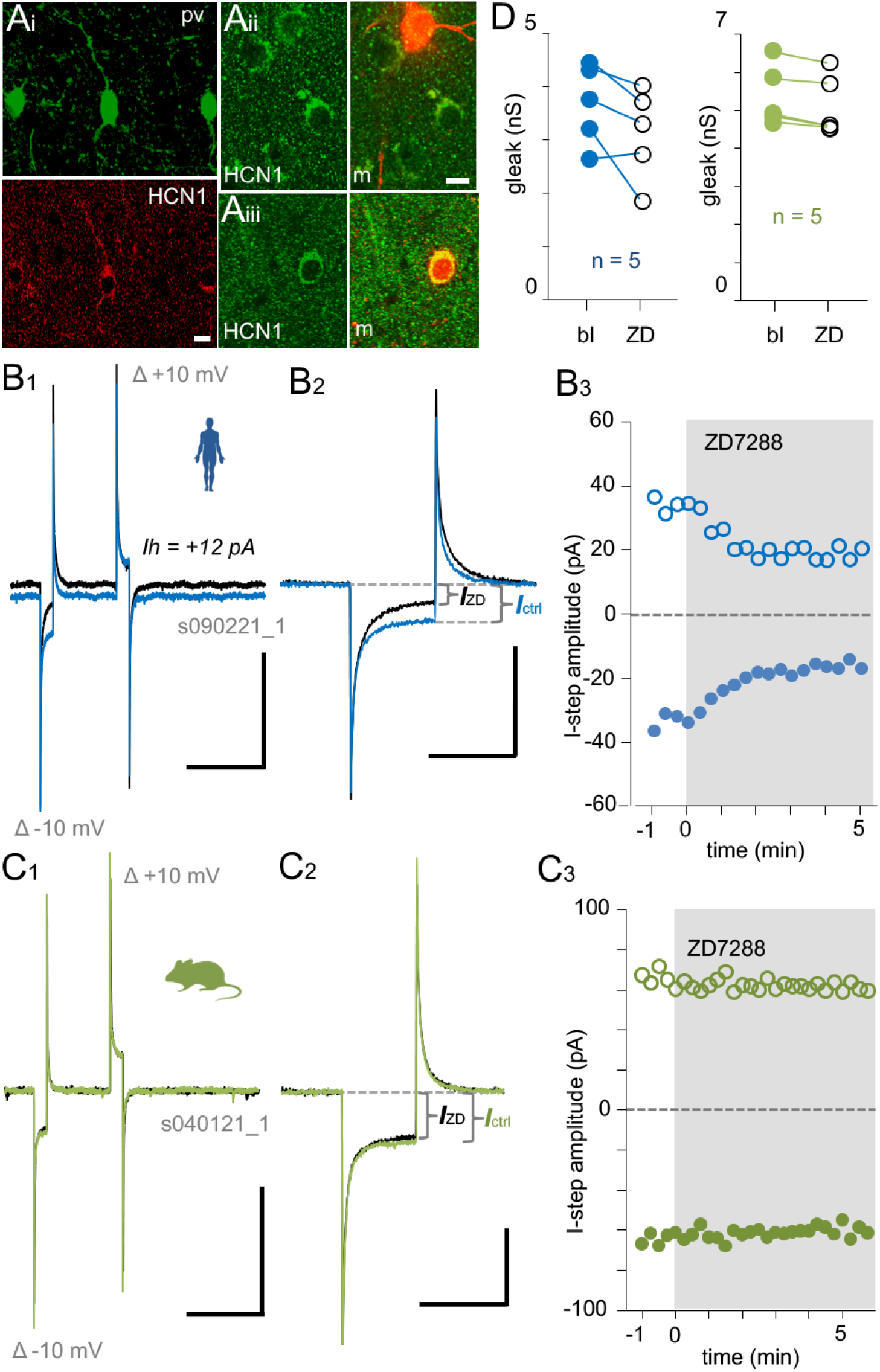
HCN channels maintain somatic leak conductance in human pv+BCs at the resting membrane potential. *A*. Human pv+ interneurons in L2/3 show strong immunopositivity for hyperpolarization-activated cyclic nucleotide-gated channel 1 (HCN1). Confocal images showing co-localization of pv and HCN1 immunofluorescence in L2/3. Micrographs Ai–iii are different neocortical samples. Alexa488 (green) and cy3 (red) (m, merge). Scale 10 μm. *B*. Voltage-clamp step shows a leak conductance reduction in human pv+BC with application of the HCN-channel blocker ZD7288 (ZD; 10 μM). *B1*. Sample traces (averages of five) show somatic voltage-clamp current for −10 and +10 mV sequential voltage steps (10 ms) measured at Em. Blue trace is in control conditions and black trace in the presence of ZD (10 μM). Voltage-clamping pv+BC at Em (−65 mV throughout the experiment) was associated with a moderate (+12 pA) shift of holding current level in ZD. Scale 100 pA, 50 ms. *B2*. Zoomed-in and superimposed traces (averages of five, the holding current levels are aligned for clarity) illustrate the voltage-clamp current to a −10 mV step in the two conditions. There is a decrease in current amplitude for the voltage step in ZD as indicated by brackets; Current step amplitude in control and in ZD are marked as I_ctrl_ and I_ZD_ respectively. I_ctrl_ or I_ZD_ amplitude is proportional to somatic leak conductance. Scale 100 pA, 10 ms. *B3*. Plot shows the current amplitude for −10 mV (solid symbols) and +10 mV (open symbols) voltage steps before and during the wash-in with ZD (indicated with gray background) in the same experiment. *C*. Voltage-clamp step current is not changed in a mouse pv+BC by ZD. *C1–3*. ZD wash-in shows negligible effect in identical experiment conducted in a mouse pv+BC. Scales: 200 pA, 50 ms and 100 pA, 10 ms. *D*. Plot compiles leak conductance (nS) values translated from the voltage-clamp step current in 5 human and 5 mouse pv+BCs in control conditions (baseline, bl) and in ZD. ZD-sensitive conductance (G_HCN_) was 11.9%, 1.8%–30% of total leak conductance in human, and 4.36%, 2.7%–5.4% in mouse cells (P = 0.151 between the human and mouse, Mann– Whitney U-test).

### HCN-channel leak conductance shortens the membrane time-constant, the EPSP time-to-peak and the EPSC-to-spike transformation in human pv+BCs

Considering the evidence presented above of HCN-channel activity in human pv+BC soma, we studied the effect of a HCN-channel blocker on somatic membrane time constant in human and mouse pv+BCs. To measure the effect of ZD (10 μM) on passive membrane time constant (membrane tau) in pv+BCs, we applied hyperpolarizing voltage steps at −70 mV (Fig. 4A1), and found that the drug wash-in was associated with a 1.33-time (average range of 1.08–1.63 in ZD compared to baseline control) prolongation of the membrane tau in human pv+BCs, and a 1.14-time (average range of 0.98–1.28) prolongation in mouse pv+BCs (Fig. 4A2) (P = 0.059, Student’s t-test). Membrane tau for human cells was 6.76 ms, 6.55– 8.22 ms in control conditions (median and quartiles, n = 8), and 9.72 ms, 8.15–10.53 ms in ZD conditions (P = 0.008, Wilcoxon Signed-Rank Test), when measured in standard 20 mV hyperpolarizing steps at −70 mV as illustrated in Figure 4A1 inset. Correspondingly, tau values in mouse pv+BCs were 5.56 ms, 5.33–6.16 ms in control and 6.95 ms, 5.40–7.42 in ZD (n = 7) (P = 0.031, Wilcoxon Signed-Rank Test). Importantly, the effect of ZD on membrane tau was most pronounced in pv+BCs showing the largest HCN voltage sag potentials in control conditions before the ZD wash-in (P > 0.001, n = 15 cells, Pearson’s correlation). Cross-correlation of the two ZD-sensitive variables is illustrated in Figure 4A3. Together, these results indicate that HCN-channel blocker affects somatic passive membrane time constant more pronouncedly in human than in mouse pv+BCs.

**Figure 4.**
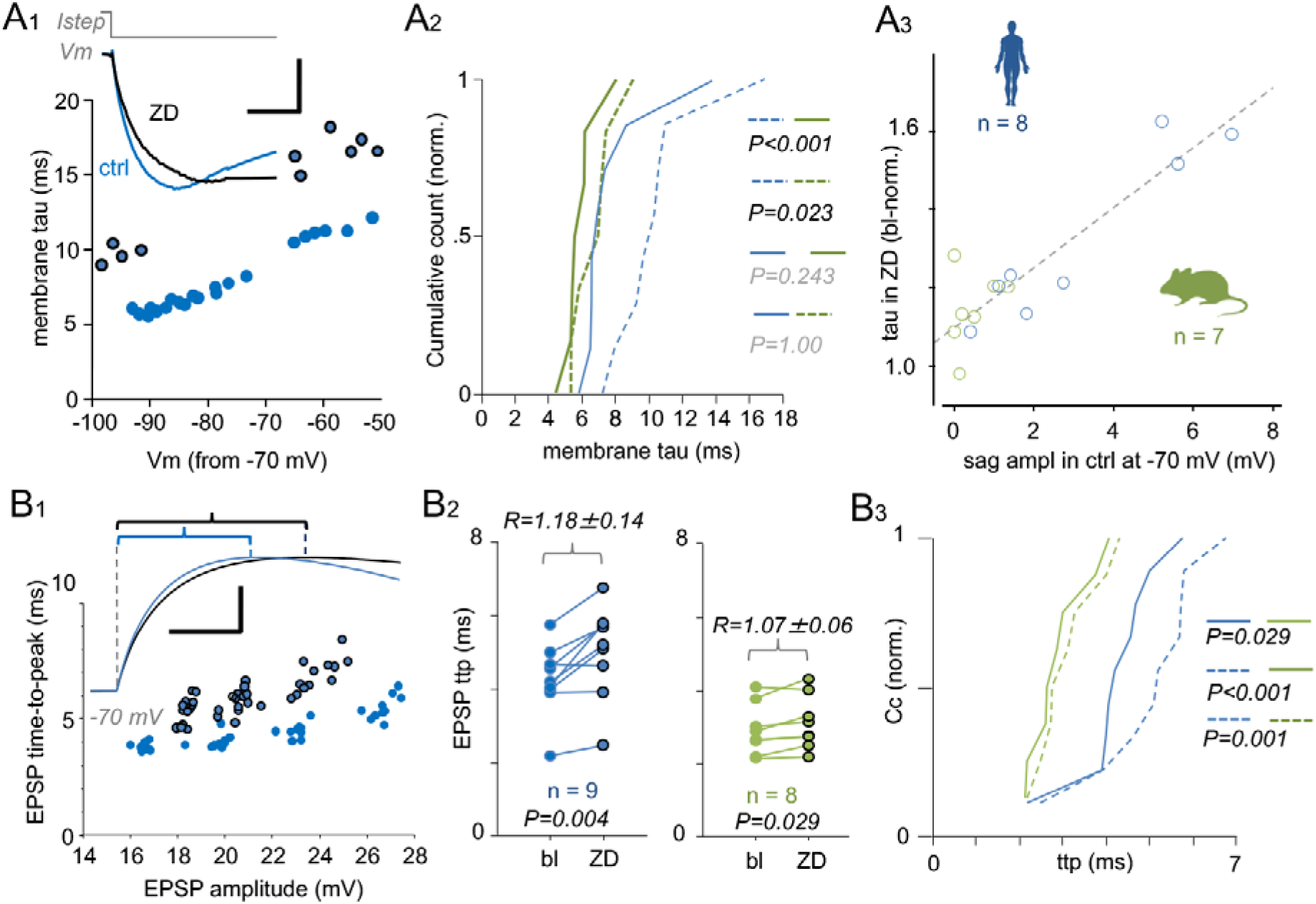
HCN-channel activity shortens the somatic time-constant and the EPSP time-to-peak in human pv+BCs. *A*. HCN-channel activity keeps human pv+BC membrane tau close to tau in mouse cells. *A1*. Plot shows membrane time constant (membrane tau) measured in human pv+BC soma during different membrane potential steps delivered at −70 mV in control conditions (blue) and in ZD7288 (ZD; black). Inset: traces illustrate membrane potential step (average of five traces) from −70 mV to −90 mV in control (blue) and in ZD (black). Scale 10 mV, 5 ms. I*step* indicates the hyperpolarizing current step. *A2*. Cumulative histograms show passive membrane time constant (membrane tau) measured during Vm steps from −70 mV to −90 mV in human (blue, n = 8) and mouse (green, n = 7) pv+BCs. Solid line is in control conditions and dotted line is in the presence of ZD. ZD significantly prolongs the passive membrane tau both in human (n = 8, P = 0.008) and in mouse (n = 7, P = 0.031) pv+BCs. Yet in the presence of ZD, human pv+BC tau (dotted blue line) is significantly different from mouse pv+BC values (green lines), whereas in control condition there is no significant difference between the human and mouse tau values (Kruskal–Wallis H-test and multiple comparisons against control group with Dunn’s method). *A3*. ZD effect on passive membrane tau is strongest in pv+BC showing the largest HCN sag potential amplitude in control conditions. Plot shows the baseline-normalized membrane tau value in ZD (ordinate) against the pv+BC HCN sag amplitude (abscissa) measured in cells under control conditions. Pearson correlation indicates linear correlation (Coefficient = 0.905, P > 0.001, n = 15 cells, r^2^ = 0.818). Dotted line shows regression. The sag and the tau were measured in Vm steps from −70 mV to −90 mV. *B*. ZD prolongs somatic EPSPs time-to-peak (applied with dynamic-clamp) in human pv+BCs. *B1*. Plot shows time-to-peak (ordinate) for different EPSP amplitudes (abscissa) in a sample human pv+BC in control conditions (blue) and in ZD (black). EPSPs were generated with incremental EPSCs strength (EPSCs not shown) at −70 mV. Inset: EPSP of similar amplitude in control and in ZD (average of five) illustrate longer time-to-peak (indicated with brackets) in the presence of ZD. Scale: 5 mV, 2 ms. *B2*. ZD effect on EPSP time-to-peak (ttp) tested in randomly selected 9 human (blue) and 8 mouse (green) pv+BCs. EPSPs were evoked by dynamic-clamp and values ttp-values are measured from EPSPs of similar amplitude in control and ZD (Human P = 0.004; mouse P = 0.029, paired t-test) (Shapiro–Wilk normality P = 0.546 and P = 0.705, respectively). EPSP ttp increases significantly both in human and in mouse cells. However, effect of ZD on EPSP time-to-peak is stronger in the human than in mouse pv+BCs (P = 0.049, t-test. Shapiro–Wilk normality P = 0.34). Baseline-normalized EPSP amplitude (R) in ZD was 1.18 ± 0.14 in human and 1.07 ± 0.06 in mouse cells. *B3*. Cumulative histogram shows the EPSP time-to-peak values in control and in ZD. EPSP ttp is longer in human than in mouse both in control conditions and in ZD. (Control, solid line; ZD dotted line). One-way ANOVA (P < 0.001) with pairwise multiple comparison (Holm–Sidak method).

By using dynamic-clamp, a computer-driven current injection system mimicking synaptic currents, we next applied somatic excitatory postsynaptic currents (EPSCs) similar to those occurring in human during pyramidal cell-to-pv+BC communication ^14, 15^, in randomly selected separate sets of pv+BC in human and mouse. In cells recorded at −70 mV, we evoked large subthreshold EPSPs nearly reaching the action potential firing threshold. The results show that HCN-channel blocker ZD prolongs the time-to-peak of the EPSPs generated by the EPSCs (Fig. 4B1). By comparing EPSPs of similar peak amplitude in the control condition and in the presence of ZD, the EPSP time-to-peak was prolonged with the drug in human (P = 0.004, n = 9) and also in mouse (P = 0.029, n = 8) cells (paired Student’s t-test) (Fig. 4B2). However, the effect of ZD on the EPSP time-to-peak was found to be larger in human compared to mouse pv+BCs (P = 0.049, Student’s t-test; Shapiro–Wilk normality P = 0.34), increasing the baseline-normalized value to 1.18 ± 0.14 in human and to 1.07 ± 0.06 in mouse pv+BCs.

Given that ZD prolongs the membrane tau and the EPSP time-to-peak in human pv+BC soma, we continued our investigation using dynamic-clamp to study the effect of HCN-channel blockade on EPSC transformation to a spike in pv+BCs. By applying incrementally increasing EPSC amplitudes (testing 4–5 different EPSC strengths) to elicit EPSPs with the peak amplitude reaching the firing threshold, we assessed the time lag for the EPSC transformation to an action potential in control conditions and after a brief wash-in (5 min) with ZD. Figure 5A illustrates the sample dynamic-clamp EPSC, the EPSP generated and the plots data on the EPSC-to-spike delay relationship in one pv+BC.

**Figure 5.**
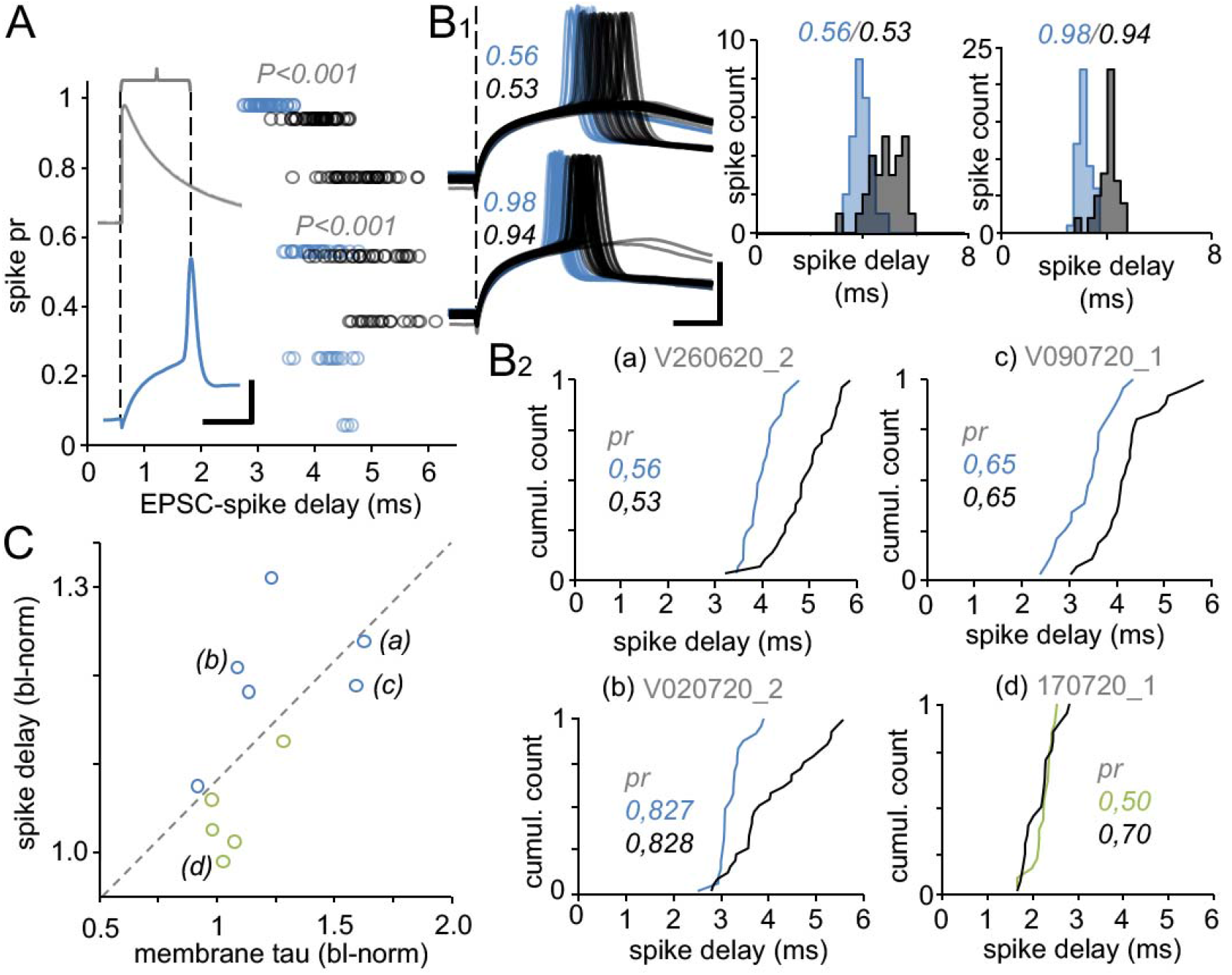
HCN-channel activity shortens EPSC-to-spike delay in human pv+BCs. *A*. Dot plot shows EPSC-to-spike delay (abscissa) in control and in the presence of ZD in one human pv+BC. Four different EPSC strength were used in both conditions to elicit EPSP-spike compound with four different spiking probability (ordinate). EPSCs are not identical strength in control and in ZD, but their strength was adjusted to aim EPSPs with similar spike probability in the two conditions. Blue and black symbols show delay of the spikes evoked when 30 cycles by each EPSC strength was tested (interval 1 s). Membrane potential is −70 mV throughout the experiment. *P* values (paired Student’s t-test) are tested for delay values when spikes were elicited with similar probability in control (blue) and in ZD (black) (control 0.56 and in ZD 0.53; control 0.98 and in ZD 0.94). Inset: traces illustrate EPSP with spike (blue) elicited with EPSC (gray, dynamic-clamp) at −70 mV. Dotted lines and the bracket illustrate EPSC-to-spike delay, with traces showing dynamic-clamp EPSC (gray) and the EPSP with spike. Scale: 200 pA, 25 mV, 2 ms. *B*. ZD increases EPSC-to-spike delay in various individual human pv+BCs. *B1*. Superimposed EPSP-spike traces in one human pv+BC (same cell as in A) in control conditions (blue) and in ZD (black). Traces show EPSP-spike compounds with spike probability of 0.5 and close to 1. Histograms summarize EPSC-to-spike delay illustrated with the traces. *B2*. Cumulative histograms show EPSC-to-spike delay in 3 human pv+BCs and in 1 mouse pv+BC under control (blue, green) and ZD (black) conditions. Spike probability value (pr for 30 EPSC cycles) is indicated in the plots. *C*. ZD effect on EPSC-to-spike delay is largest in pv+BCs showing the strongest effect of ZD on the passive membrane time-constant (measured during the hyperpolarizing step). Pearson correlation indicates linear correlation (Coefficient = 0.619, P = 0.0129, n = 11 cells, r^2^ = 0.383). Blue, human pv+BCs, green, mouse pv+BCs. Dotted line shows regression.

Because ZD increases EPSP amplitude by EPSCs, but not the relation of EPSP amplitude and spiking probability (Supplementary Fig. S2–S3), we compared the spike delay between similar amplitude EPSPs in the control and ZD conditions. Figure 5B1 illustrates one experiment in human pv+BCs with EPSP-spike probability close to 0.5. In the presence of ZD, we set EPSC strength to evoke EPSP with spiking at a similar probability as in the control condition. The probability was set to 0.5 < 1. To eliminate Em shift caused by ZD, and thus, to focus on the HCN-channel effect on the somatic membrane time constant and lag in the spike generation, the membrane potential was held at −70 mV throughout the experiments. We found that the EPSC-to-spike delay increased in five out of six human pv+BCs but only in one out of five mouse pv+BC. The results are illustrated in Figures 5B2-C. Importantly, we compared the spike-delay in ZD (normalized by spike delay in control) with the cell’s passive membrane time constant in ZD (normalized by the value in control conditions), revealing a strong correlation between them (P = 0.013, Pearson’s test) (Fig. 5C).

### HCN-channel conductance in soma facilitates pv+BC input-to-output function

Finally, we investigated the effect of somatic HCN conductance on pv+BCs in a computational model. We used a single-cell soma model (NEURON-7) with passive membrane properties to simulate the effect of G_HCN_ on the cell Em, the EPSP amplitude and the EPSP time-to-peak at the same time, since these features were studied separately in the experiments above. We simulated the five human and five mouse pv+BCs with cell capacitance (Cm), Em measured in control conditions, total leaking conductance measured in control conditions, and the ZD-sensitive leaking conductance (measured in the cells presented in Fig. 3 and summarized in Supplementary Table 3). To elicit the EPSPs, we used same EPSC parameters as in the dynamic-clamp experiments above (see Figs. 4–5). The simulation demonstrated a strong faciliatory effect of the G_HCN_ in human pv+BCs, with the conductance values occurring naturally in these cells (Fig. 6A). Importantly, the simulations demonstrate that although the somatic HCN conductance leads to an attenuated EPSP amplitude in pv+BCs, the simultaneous depolarization of Em by somatic G_HCN_-mediated depolarizing leaking current (with a reversal potential at −30 mV) ^40, 42^ overwhelms the EPSP amplitude attenuation. Thus, in all cells with natural G_HCN_ measured in their soma, this conductance promotes EPSC-to-spike transformation; it allows for a somatic excitatory current to reach higher depolarization level from Em, and in addition it facilitates the EPSP time-to-peak to reach any depolarized threshold level with a shorter delay. Figure 6A1-3 illustrates the effects of G_HCN_ in three human pv+BCs. Correspondingly, ZD-sensitive conductance measured in mouse pv+BC soma had no effect or only a small effect on cell Em or EPSP. Figure 6B1-3 shows the simulated EPSPs at Em in three mouse pv+BCs. Thus, the simulations demonstrate that the net effect of somatic HCN conductance is a facilitation of the excitatory input-to-output function (Fig. 6C1-3), which speeds up the EPSC-EPSP transformation to reach firing threshold in these cells (Fig. 6D).

**Figure 6.**
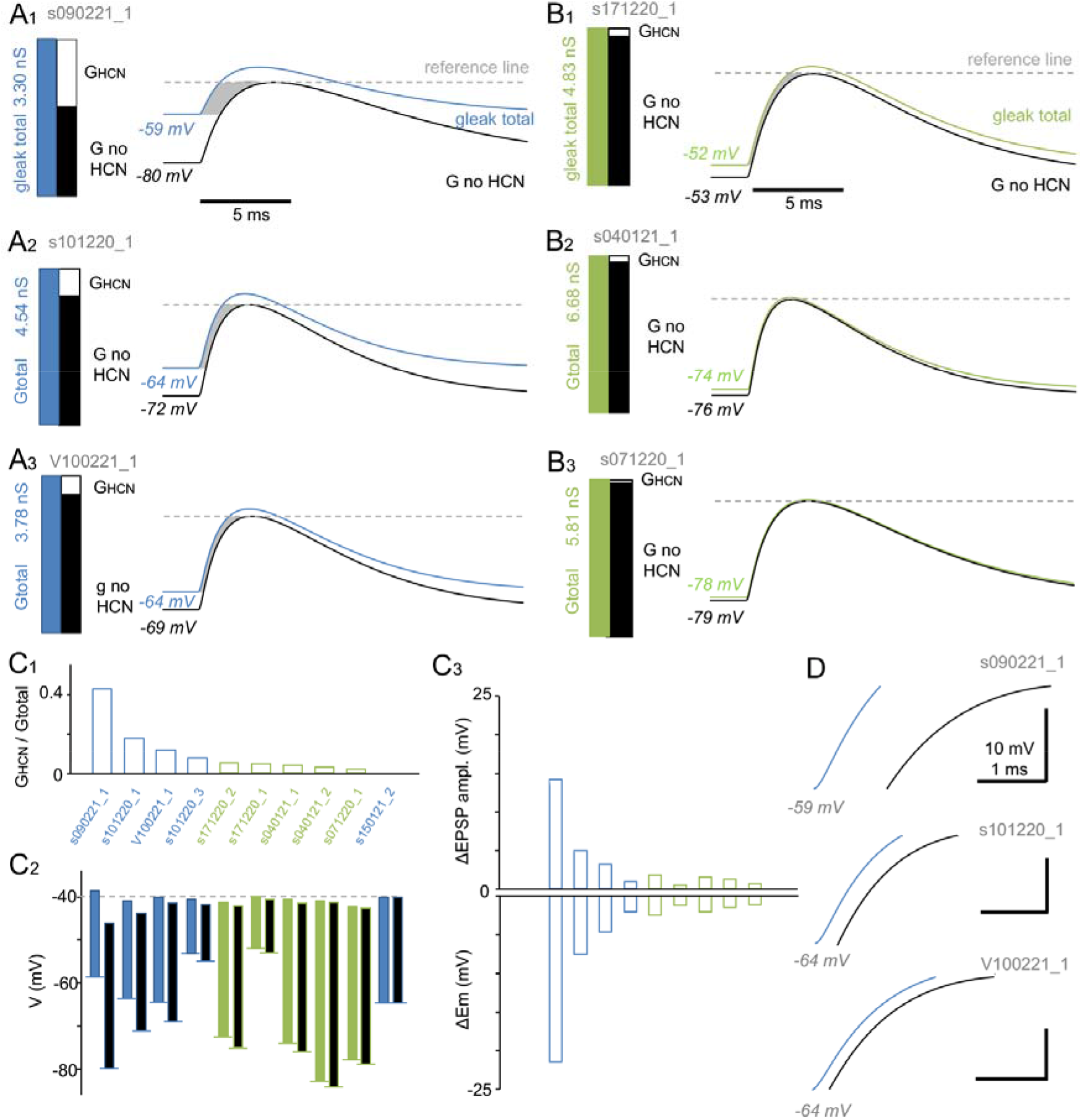
Somatic HCN conductance facilitates pv+BC input-to-output function in a computational model. HCN channel-mediated leaking conductance (G_HCN_) in the human pv+BCs attenuates somatic EPSP amplitude. However, a large depolarizing effect of G_HCN_ on the resting membrane potential (Em) dominates the EPSP amplitude attenuation. The net effect by G_HCN_ is that EPSCs generate EPSPs reaching higher depolarizing level and with shorter delay. *A*. Computational single-cell passive membrane model replicates human pv+BCs with authentic cell capacitance, Em, G_HCN_, and ZD-insensitive leak conductance measured in the cells. The analysis includes 5 human and 5 mouse pv+BC studied in Figure 3. *A1–3*. Illustration of three human pv+BC simulations. Traces show EPSPs evoked in the soma by same EPSC (3.4–5.0 nS) at resting membrane potential (Em) in control conditions (blue) or without the ZD-sensitive G_HCN_ (Erev = −30 mV). Bar charts show the relative proportion of G_HCN_ (white area) and the ZD-insensitive conductance (g no HCN, black area) of the total leak conductance (blue bar) measured in the cell in control conditions. In the absence of G_HCN_, Em is hyperpolarized (black font). Horizontal dotted line shows a reference line to compare the membrane potential depolarization level reached by EPSPs at specific time point. Shaded gray area highlights the time delay between the EPSPs to reach a nominal depolarization level. *B*. Simulation experiments illustrated in three mouse cells (*B1–B3*) shows that the modest somatic G_HCN_ in these cells has negligible effect on somatic Em or the EPSPs. *C*. Modeling data demonstrate that net effect of somatic G_HCN_ in pv+BCs is their facilitated excitation. *C1*. Bar histogram shows the ratio of G_HCN_ to the total leak conductance (Gtotal) measured in the 5 human (blue) and 5 mouse (green) pv+BCs at Em. Abscissa shows the cell codes. *C2*. Same cells as in C1 showing the EPSP amplitude (vertical bar) at Em (horizontal line at bottom of the bars) in control conditions (with G_HCN_, blue or green) and in the absence of G_HCN_ (black). EPSC underlying the EPSPs differs in strength between the cells, and it was adjusted to elicit an EPSP that reaches the average firing threshold level (indicated by horizontal dotted line) in the control conditions. The bars show that with G_HCN_ in cell soma, the Em depolarizes more than the EPSP amplitude attenuates. When G_HCN_ is off, the EPSPs (black bars) fail to reach the firing threshold level from the hyperpolarized Em. *C3*. Summary of the G_HCN_ effect on EPSP amplitude and Em in the pv+BCs. Vertical bars demonstrate the increase of EPSP amplitude (upper bars) and the negative shift of Em (lower bars) in the absence of G_HCN_ in the cells. The shift in Em is larger than the increase in EPSP amplitude (n = 10, P = 0.002, Wilcoxon Signed-Rank Test). *D*. Zoomed-in traces showing the shaded area between EPSPs in the 3 human pv+BCs illustrated in *A1-3*. Depolarization by EPSPs is slower in the absence of G_HCN_ (black).

## DISCUSSION

Our results reveal robust somatic HCN-channel activity in pv+BCs of the human neocortex. The somatic HCN-channel responses are absent in pv+BCs of the mouse neocortex, as was reported in various earlier studies ^33, 36, 37, 38, 39, 43, 50^. These findings demonstrate, together with the computational modeling, how “a human-specific” physiological feature augments and speeds up signal transfer in the highly abundant inhibitory circuit in neocortex. This emphasizes the importance of conducting physiological studies in human identified cell types in order to uncover their “uniqueness.”

### Between-species difference in somatic HCN activity alters basket cell physiological function

HCN channels in the somatic compartment modulate the entire human pv+BC function through the leaking conductance at potentials more negative than −50 mV. The HCN-mediated somatic conductance facilitates their EPSP rise, and although it reduces the EPSP peak amplitude, it depolarizes the resting potential more than it reduces the EPSP amplitude. A similar net effect of HCN leak conductance was reported in glutamatergic neurons in rodent and human ^18, 52, 53^. We show here that the somatic HCN facilitates input- to-output function in the most abundant neocortical interneuron type in human but not mouse.

The somatic membrane time-constant is a key determinant of the input-output relation of neurons and it determines how rapidly a neuron responds to an excitatory stimulus ^52^. In the human neocortex, unitary pyramidal cells form powerful excitatory connections on local pv+BCs, in order to generate suprathreshold EPSPs at the interneuron resting potential by unitary spike ^10, 14, 15, 21^. In this input-to-output signaling, the pv+BC’s firing delay to the large EPSC is couple of milliseconds ^10, 12, 21 54^. In the rapid EPSP-spike transformation, the somatic G_HCN_ at Em is highly relevant, considering that HCN-channel conductance remains practically unchanged during fast Vm shifts such as EPSP rise and the EPSP-spike coupling. Although G_HCN_ depends on the membrane potential, it changes relatively slowly by membrane potential shifts with a time constant of 20–50 ms ^40, 55^.

Together, our results demonstrate that with the contribution of somatic HCN, human pv+BCs keep their time-constant and EPSP time-to-peak kinetics to a similar level as in rodent cells. Without the somatic G_HCN_, human cells would be substantially slower than their rodent counterparts. Thus, G_HCN_ activity appears critical for the rapid membrane potential dynamics in human pv+BCs.

### Rationale of basket cell specialization is efficiency

Although benefits of somatic G_HCN_ in neuronal signaling are obvious, a question remains: why has this mechanism evolved in human (and presumable in other primate) neocortical pv+BCs but not in the same cell type in rodent neocortex?

Firstly, we speculate that in general there is evolutionary pressure toward faster and higher fidelity signaling in neuronal pathways ^22^. Secondly, many human neurons including the pv+BCs exhibit highly resistive cell membrane and a relatively large cell capacitance, which together result in slow passive membrane time constant and EPSP-to-spike transformation in cell soma ^19, 28, 31^. This was observed here in the experiments blocking HCN channels. Thus, the active leak conductance from somatic G_HCN_ channels appears critical in human cells to speed up their membrane potential dynamics. For some unknown reason, the phylogenetic development of BC intrinsic properties has followed a different path in the rodent neocortex, since these cells lack or show very small somatic G_HCN_ although they have prominent somatic leaking conductance ^33, 36, 43^. Apparently, rodent pv+BCs maintain their somatic leakiness (and reduce their membrane time constant) through K^+^ channels and voltage-gated Cl^−^ channel conductance instead ^4, 56^. Interestingly, like human pv+BCs, fast-spiking neocortical basket cells of macaque monkeys exhibit high input resistance, a relatively long passive membrane time constant, and a large HCN channel-mediated voltage sag, suggesting robust somatic HCN-channel activity ^33, 37, 50^. Thus, somatic HCN-channel leak conductance in neocortical pv+BCs may be a general feature encountered across primates ^57^ which separated from rodents 70–90 million years ago^58^.

Overall, these results support the need for more functional studies with quantitative analysis of the physiological parameters of identified human neuron types. This knowledge is required to uncover the human “uniqueness” across various neuroanatomical circuits. By employing computational models, the generated data will help us to understand why species–specific neuronal features are important and relevant to cortical processes. Some of these features may eventually turn out to be key targets for neurological and neuropsychiatric disorders as well as neurodegenerative diseases ^16, 59, 60, 61, 62^.

## METHODS

### Ethics statement

All the procedures were performed according to the Declaration of Helsinki with the approval of the University of Szeged Ethical Committee and the Regional Human Investigation Review Board (ref. 75/2014). For all human tissue material, written consent was obtained from patients prior to surgery. Tissue obtained from minor patients was provided with agreement from a parent or legal guardian.

### Human brain slices

Neocortical slices were sectioned from material that had to be removed to gain access for the surgical treatment of deep-brain targets from the frontal, temporal, or occipital areas. The patients were 14–74 years of age, and samples from males and females from either the left or right hemisphere were included. Anesthesia was induced with intravenous midazolam and fentanyl (0.03 mg/kg, 1–2 lg/kg, respectively). A bolus dose of propofol (1–2 mg/kg) was administered intravenously. The patients also received 0.5 mg/kg rocuronium to facilitate endotracheal intubation. The trachea was intubated, and patients were ventilated with an O_2_/N_2_O mixture at a ratio of 1:2. Anesthesia was maintained with sevoflurane. Following surgery, the resected tissue blocks were immediately immersed into ice-cold solution within a glass inside the operating room. The solution contained (in mM): 130 NaCl, 3.5 KCl, 1 NaH_2_PO_4_, 24 NaHCO_3_, 1 CaCl_2_, 3 MgSO_4_, 10 D(+)-glucose and was saturated with 95% O_2_/5% CO_2_. The container was placed on ice in a thermally isolated transportation box where the liquid was continuously gassed with 95% O_2_/5% CO_2_. Then, the tissue was immediately transported from the operating room to the electrophysiology laboratory where slices of 350 μm thickness were prepared from the tissue block with a vibrating blade microtome (Microm HM 650 V). The slices were incubated at 22°C–24°C for 1 h, when the slicing solution was gradually replaced by the solution used for storage (180 ml) using a pump (6 ml/min). The storage solution was identical to the slicing solution, except that it contained 3 mM CaCl_2_ and 1.5 mM MgSO_4_.

### Drug

ZD7288 (Sigma Aldrich, Budapest, Hungary) was diluted in physiological extracellular solution and applied by wash-in.

### Identification of fast-spiking BCs in mouse brain slices

Transversal slices (350 μm) from the somatosensory and in some cases the frontal cortex were prepared from 4- to 6-week-old heterozygous B6.129P2-Pvalbtm1(cre)Arbr/J mice (The Jackson Laboratory, stock 017320, B6 PVcre line) expressing td-Tomato fluorophore preferentially in parvalbumin GABAergic neurons. Cells were confirmed to be fast-spiking by electrophysiological recording of fast spike kinetics and high-frequency non-accommodating firing pattern to suprathreshold depolarizing 500 ms pulses. Cells were visualized with streptavidin Alexa488 (1:2000, Jackson ImmunoResearch Lab, Inc.) and identified visually under epifluorescence microscopy.

### Electrophysiology

Recordings were performed in a submerged chamber (perfused 8 ml/min) at 36°C–37°C. Cells were patched using a water-immersion 20x objective with additional zoom (up to 4x) and infrared differential interference contrast video microscopy. Micropipettes (5–8 MΩ) for whole-cell patch-clamp recording were filled with intracellular solution (in mM): 126 K-gluconate, 8 NaCl, 4 ATP-Mg, 0.3 Na_2_–GTP, 10 HEPES, and 10 phosphocreatine (pH 7.0–7.2; 300 mOsm) with 0.3% (w/v) biocytin. Recordings were performed with a Multiclamp 700B amplifier (Axon Instruments) and low-pass filtered at a 6–8 kHz cut-off frequency (Bessel filter). Series resistance and pipette capacitance were compensated in current-clamp mode and pipette capacitance was compensated in voltage-clamp mode. Liquid junction potential error was not corrected in membrane potential values. The access resistance of the recording electrode was measured, and its effect on the clamping potential error was corrected in nominal somatic potential reading. Em was recorded 1–3 min after break-in to whole cell. Cell capacitance was measured in current clamp using −50–100 pA, 200–300 ms steps delivered at Em. The somatic leaking conductance in Figure 3 was measured in voltage clamp at Em by delivering −10 mV and 10 mV (10 ms) voltage steps, and conductance was calculated from clamping current amplitude as demonstrated in Figure 3 using the Ohm’s equation. Somatic input resistance in current clamp was measured from hyperpolarizing (−10 to −20 mV, 250 ms) voltage steps at −70 mV.

### Dynamic clamp

To induce EPSPs, a software-based dynamic-clamp system was employed. Current injections were calculated and delivered by the Signal software (Cambridge Electronic Design, Cambridge, UK) through a Power1401-3A data acquisition interface (CED, Cambridge, UK) based on voltage signals of the electrode. We ran the dynamic clamp on a computer distinct from our experimental data acquisition system (recording cell membrane potential) to record and verify dynamic-clamp output (conductance and EPSCs). EPSCs were generated using a decay time constant of 3 ms and a reversal potential of 0 mV. The peak conductance (1.5–10 nS) was set to evoke EPSPs.

### Single-cell model

For the simulation of somatic EPSP, real experimental data from individual basket cells were used. Simulation of a basket cell membrane potential was performed using a NEURON 7.6.5 simulator (Carnevale NT HMC, UK: Cambridge University Press 2006). The membrane capacitance was 1 μF/cm^2^, and the size of the soma was determined so that the total cell capacitance matched the actual measured value in each cell. G_leak_ (total G at resting Em) and G_HCN_ (ZD-sensitive conductance) were retrieved from voltage-clamp data in Figure 3 human and mouse identified neurons. G_HCN_ reversal potential was −30 mV, and G_leak_ reversal potential was set at the resting membrane potential value measured in the presence of ZD in each recorded cell to reach the actual Em value measured in control conditions. EPSPs were modeled using a rise tau of 0.2 ms, decay tau of 3 ms and conductance of 10 nS with a reversal potential of 0 mV ^12, 14^.

### Data analysis

Data were acquired using the Clampex software (Axon Instruments) and digitized at 35–50 kHz. The data were analyzed off-line with pClamp (Axon Instruments), Spike2 (version 8.1, Cambridge Electronic Design), OriginPro (OriginLab Corporation) and SigmaPlot14 software (Systat Software).

### Statistics

Data are presented as the mean ± SD, for samples with size (n) ≥ 7 and a parametric distribution. Normality was tested with the Shapiro–Wilk test using a P value > 0.05. Otherwise, data are shown as the median with interquartile range (of lower and upper quartile) or as the average and range, unless stated otherwise. Correspondingly, for statistical analysis, Student’s two-tailed or paired t-test, Mann–Whitney U-test, Wilcoxon Signed-Rank Test or two-tailed ANOVA or Kruskal-Wallis H-test (with Dunn’s *post-hoc* test) were used with Sigma Plot. Correlations were tested using Pearson or Spearman correlation, respectively. Differences were considered significant at P < 0.05.

### Tissue fixation and cell visualization

Biocytin-filled cells were visualized by using either Alexa488- (1:500) or Cy3-streptavidin (1:400, Jackson ImmunoResearch Lab, Inc.). After electrophysiological recording, slices were immediately fixed in a fixative solution containing 4% paraformaldehyde and 15% picric acid in 0.1 M phosphate buffer (PB, pH = 7.4) at 4°C for at least 12 h and then stored at 4°C in 0.1 M PB containing 0.05% sodium azide as a preservative. All slices were embedded in 10 % gelatin and further sectioned into slices of 60 μm thickness in ice-cold PB using a vibratome VT1000S (Leica Microsystems). After sectioning, the slices were rinsed in 0.1 M PB (3 × 10 min) and cryoprotected in 10–20 % sucrose solution in 0.1⍰M PB. Afterwards, the slices were flash-frozen in liquid nitrogen and thawed in 0.1⍰M PB. Finally, they were incubated in a fluorophore-conjugated streptavidin (1:400 or 1:500, Jackson ImmunoResearch Lab, Inc.) in 0.1 M Tris-buffered saline (TBS, pH 7.4) for 2.5 h (at 22°C–24°C). After washing with 0.1 M PB (3 × 10 min), the sections were covered in Vectashield mounting medium (Vector Laboratories Inc.), placed under coverslips, and examined under an epifluorescence microscope with 20-60 × magnification (Leica DM 5000 B).

### Cell reconstruction and anatomical analyses

Sections used for morphological analyses in Figure 1C were further incubated in a solution of conjugated avidin-biotin horseradish peroxidase (ABC; 1:300; Vector Labs) in TBS (pH = 7.4) at 4°C overnight. The enzyme reaction was revealed with the glucose oxidase-DAB-nickel method using 3⍰3-diaminobenzidine tetrahydrochloride (0.05 %) as the chromogen and 0.01% H_2_O_2_ as the oxidant. Sections were further treated with 1 % OsO_4_ in 0.1 M PB. After several washes in distilled water, sections were stained in 1 % uranyl acetate and dehydrated in an ascending series of ethanol concentrations. Sections were infiltrated with epoxy resin (Durcupan) overnight and embedded on glass slides. For the cell shown in the Figure 1 three-dimensional light microscopic reconstructions from two sections were carried out using the Neurolucida system with 100x objective (Olympus BX51, Olympus UPlanFI). Images were collapsed in the z-axis for illustration. For laminar depth analysis shown in Figure 1, the soma location and the borders of L1/L2 and L3/L4 were defined in individual 60 μm-thick sections using a brightfield microscope following the conversion to DAB.

### Immunohistochemistry

Free-floating sections were washed 3 times in TBS-TritonX 0.3 % (15 min) at 22°C–24°C and then transferred to 20 % blocking solution with horse serum in TBS-TritonX, 0.3 % for parvalbumin (pv) staining. For some sections, pre-treatment with pepsin was performed to improve immunohistochemical staining. The sections were treated with 1 mg/ml pepsin (catalog #S3002; Dako) in 1 M HCl with 0.1 M PB at 37°C for 6 min and then washed in 0.1 M PB. All sections were incubated in primary antibodies diluted in 1 % serum in TBS-TritoX 0.3 % over three nights at 4°C, and then placed in relevant fluorochrome-conjugated secondary antibody solutions (1 % blocking serum in TBS-TritonX 0.3 %) overnight at 4°C. Sections were first washed in TBS-TritonX 0.3 % (3 × 20 min) and later in 0.1 M PB (3 × 20 min) and mounted on glass slides with Vectashield mounting medium (Vector Lab, Inc.). The characterizations of primary antibodies used in humans: mouse anti-pv (1:500, Swant, Switzerland, www.swant.com, clone: 235); rabbit anti-HCN1 1: 500, MyBioSource). Fluorophore-labeled secondary antibodies were (DAM DyLight 488 donkey anti-mouse, 1:400, Jackson ImmunoResearch Lab. Inc., https://www.jacksonimmuno.com) or (DAM Cy3 donkey anti-mouse, 1:400, Jackson ImmunoResearch Lab. Inc., https://www.jacksonimmuno.com) and (DARb Cy5 donkey anti-rabbit, 1:500, Jackson ImmunoResearch Lab. Inc.). The immunoreactions were evaluated using first epifluorescence (Leica DM 5000 B) and then laser scanning confocal microscopy (Zeiss LSM880). All the micrographs presented are confocal fluorescence microscopy images.

---

**Supplementary Figure S1.**
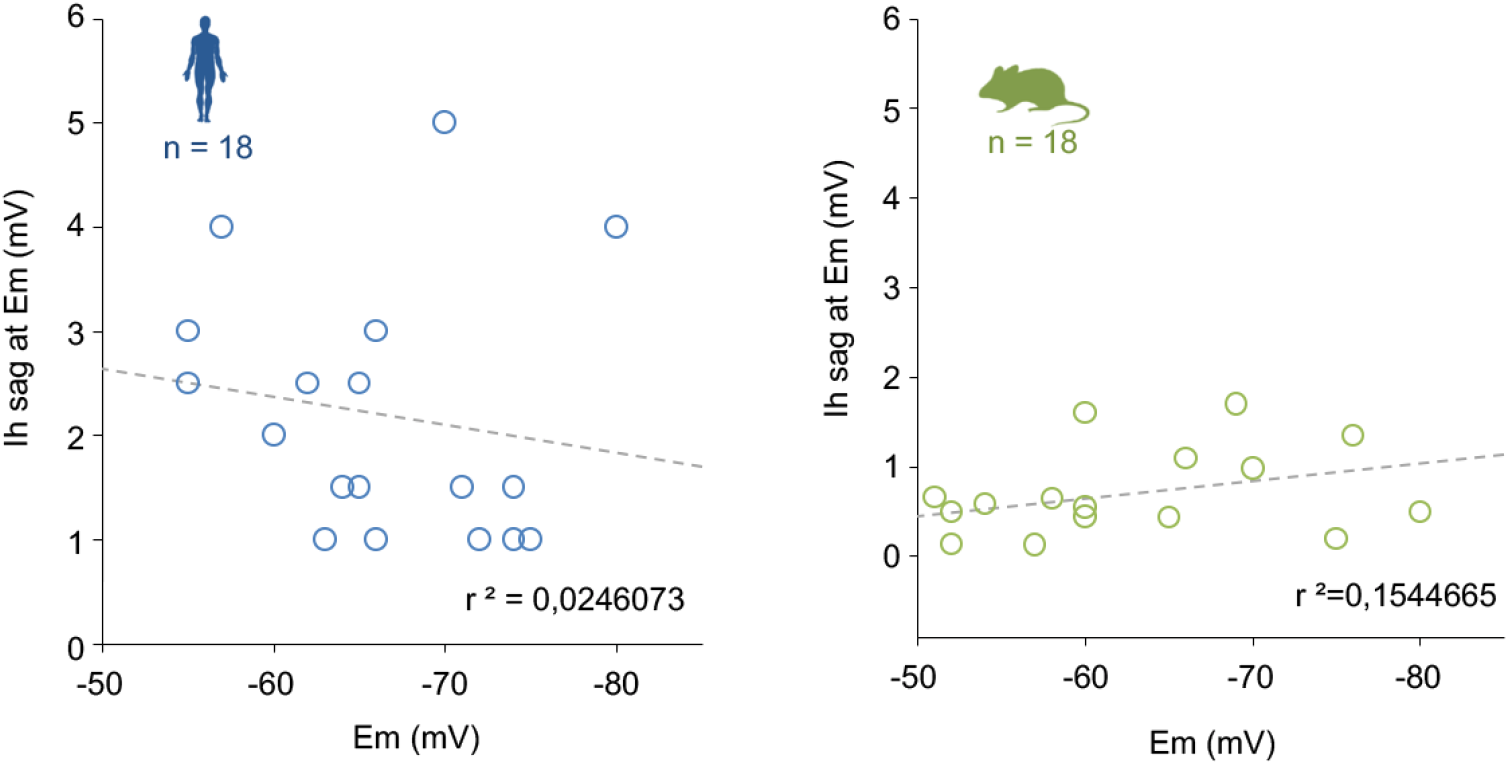
HCN potential alone does not explain Em variation in pv+BCs. HCN sag potential amplitude (measured at Em) does not show correlation with the Em value in either human (correlation coefficient = 0.314, P = 0.198) or mouse (0.416 and 0.084, respectively) pv+BCs (spearman Rank order correlation). Dotted line indicates regression.

**Supplementary Figure S2.**
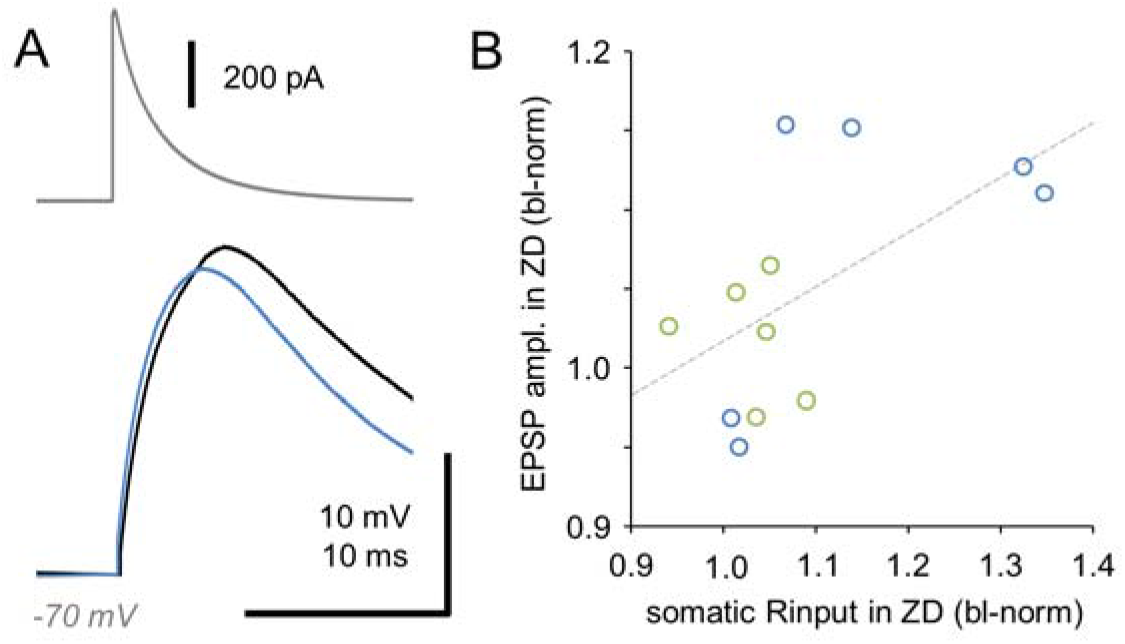
Blocking G_HCN_ increases somatic EPSP amplitude. A. Dynamic-clamp EPSC (gray trace) in a sample recording from human pv+BC demonstrates increased EPSP amplitude in the presence of ZD (black trace) compared to baseline control (blue trace). B. The plot shows that the increase in amplitude correlates with an increase of the somatic passive input resistance Rm (measured from a hyperpolarizing −20 mV step at −70 mV in current clamp) (P = 0.037, Spearman Rank Order Correlation. r^2^ = 0.327). The EPSP amplitude increase by ZD effect is smaller than expected from the changes in passive membrane properties, probably because robust depolarizing EPSPs close to firing threshold are additionally associated with voltage-sensitive conductances. Data include 6 human and 4 mouse pv+BCs.

**Supplementary Figure S3.**
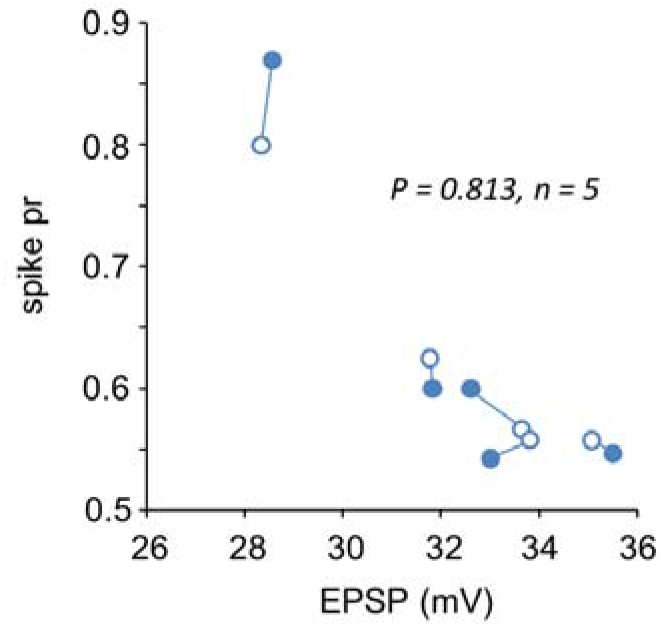
Spiking probability evoked by equal-amplitude EPSP is not significantly different between baseline control and ZD. Spiking probability evoked by equal-amplitude EPSPs (dynamic-clamp) delivered from −70 mV in baseline control (open symbols) and in the presence of ZD (solid symbols) (human, left; mouse, right). Probability of spiking (P), evoked by EPSPs applied at −70 mV in baseline and following wash-in of ZD is not changed (Wilcoxon Signed-Rank Test P = 0.688, n = 7).

Supplementary Table 1. List of anatomically identified and pv+ immunopositive human basket cells, and anatomically identified mouse fast-spiking basket cells of the study. Sheets show codes of the experiments included in specific datasets and figures.

Supplementary Table 2. A Voltage sag (‘sag during’) and a rebound sag (‘rebound sag at Em) amplitude in individual experiments (identified by cell code). ‘Step from Em’ shows the hyperpolarizing step peak amplitude (step delivered at −70 mV).

Supplementary Table 3. Table shows intrinsic pvBC parameters used for modelling in Figure 6.

## ACKNOWLEDGEMENTS

This work was supported by OTKA K 134279 (VS, SF, KL), the National Brain Research Program Hungary (VS, GT and KL), the European Research Council INTERIMPACT project (GT), the Hungarian Academy of Sciences (GT), the National Research, Development and Innovation Office of Hungary (GINOP-2.3.2-15-2016-00018, VKSZ-14-1-2015-0155, GT), the Ministry of Human Capacities, Hungary (grant 20391-3/2018/FEKUSTRAT, GT) and by University of Szeged Open Access Fund (Grant number: 4373). We acknowledge Ms Leona Mezei and Drs Gábor Molnár and Katalin Kocsis for technical assistance.

